# Phylogeographical evidence for historical long-distance dispersal in the flightless stick insect *Ramulus mikado*

**DOI:** 10.1101/2023.05.15.540668

**Authors:** Kenji Suetsugu, Tomonari Nozaki, Shun K Hirota, Shoichi Funaki, Katsura Ito, Yuji Isagi, Yoshihisa Suyama, Shingo Kaneko

## Abstract

Exploring how organisms overcome geographical barriers to dispersal is a fundamental question in biology. Passive long-distance dispersal events, although infrequent and unpredictable, have a considerable impact on species range expansions. Despite limited active dispersal capabilities, many stick insect species have vast geographical ranges, indicating that passive long-distance dispersal is vital for their distribution. A potential mode of passive dispersal in stick insects is via the egg stage within avian digestive tracts, as suggested by experimental evidence. However, detecting such events under natural conditions is challenging due to their rarity. To indirectly assess the importance of historical avian-mediated dispersal, we examined the population genetic structure of the flightless stick insect *Ramulus mikado* based on a multifaceted molecular approach (COI haplotypes, nuclear SSR markers, and genome-wide SNPs). Subsequently, we identified unique phylogeographic patterns, including the discovery of identical COI genotypes spanning considerable distances, which substantiates the notion of passive long-distance genotypic dispersal. Overall, all the molecular data revealed low and mostly non-significant genetic differentiation among populations, with identical or very similar genotypes across distant populations. We propose that long-distance dispersal facilitated by birds is the most plausible explanation for the unique phylogeographic pattern observed in this flightless stick insect.

## 1. Introduction

The segregation of populations by physical barriers and their dispersal across such obstacles constitute two prominent antagonistic forces shaping the distribution and speciation of organisms [1]. The development of wings in insects is widely regarded as a significant contributor to their prosperity and diversity [2,3], as it has facilitated predator evasion, prey capture, and migration [2]. Nevertheless, the loss or reduction of wings in various insect lineages is well documented [4], exerting a profound influence on the biogeographical and speciation patterns of these lineages [5,6]. The loss of flight within a species can restrict dispersal capabilities and foster genetic differentiation among populations, potentially leading to an elevated speciation rate in flightless lineages [5], while the evolution of flight in insects played a crucial role in their early diversification [7].

Consequently, flightless insects offer intriguing models for examining population genetic structure, with numerous species demonstrating substantial genetic differentiation between populations within relatively small geographic distances [5,6]. Phasmatodea (stick insects), encompassing over 3,000 extant species of terrestrial herbivores, primarily possess a tropical and subtropical distribution and largely consist of flightless species [8]. Phasmids are recognized for their limited dispersal capacity, with approximately 60% of all phasmid species either displaying significantly reduced wings or entirely lacking wings in their adult form [9]. Furthermore, phasmids with wings may still be incapable of sustained flight [10]. Stick insects have evolved several pivotal adaptations to counterbalance their loss of motility, such as masquerade crypsis and parthenogenesis [11]. The transition from sexual reproduction to parthenogenesis, a form of asexual reproduction, might be correlated with the flightless nature of stick insects, which renders locating mating partners more challenging [11].

Stick insects display remarkable masquerade crypsis as a defensive mechanism against visually hunting avian predators, by morphologically and behaviorally imitating twigs, bark, lichen, moss, and leaves [11,12]. Additionally, stick insects utilize camouflage techniques in their eggs, which often closely resemble plant seeds. Fascinatingly, the eggs of some stick insect species not only imitate seed appearances but also employ analogous dispersal mechanisms. Numerous stick insect eggs feature a specialized knob-like structure known as a capitulum, which closely resembles the elaiosome of ant-dispersed seeds in both form and function and shares a similar chemical composition [13]. It is hypothesized that both elaiosomes and capitula, being lipid-rich, have evolved to promote ant-mediated dispersal. Furthermore, the considerably hardened egg capsule in Euphasmatodea (including all phasmids except *Timema*) is regarded as a crucial innovation in phasmid evolution [11]. This allows the eggs to endure potentially damaging falls from the canopy and remain buoyant on seawater for extended periods [11,14].

The sturdiness of phasmid eggs might allow them to remain viable even when contained within gravid female stick insects consumed by avian predators [12,15]. This mechanism is unattainable for many other insect species, as they generally fertilize their eggs immediately before oviposition, using sperm stored in the female’s seminal vesicle after copulation. Conversely, numerous stick insects exhibit parthenogenesis, allowing them to produce viable eggs without fertilization [16]. In such instances, predation on gravid female stick insects could facilitate offspring dispersal, similar to the internal seed dispersal seen in plants producing fleshy fruits consumed by frugivorous birds. Nonetheless, it is vital to acknowledge that stick insects have developed a cryptic appearance as a means to avoid predation, rather than actively attracting animals [12]. Furthermore, we note that the study was based on a laboratory feeding experiment, necessitating caution in generalizing the findings to wild populations.

Due to their infrequent occurrences, it is likely challenging to demonstrate that avian predation serves as a factor promoting dispersion in natural settings [1]. One method to indirectly evaluate the importance of historical avian-mediated dispersal involves analyzing the spatial genetic structures of species with limited mobility, which would otherwise exhibit significant genetic differentiation among populations [1,17,18]. For example, mitochondrial DNA genetic comparisons have implied a minimum of two successful dispersal events between the Pacific and the Atlantic, traversing the digestive systems of birds, as the most plausible explanation for the long-distance dispersal of a certain marine snail [1]. Correspondingly, other investigations have proposed that long-distance dispersal facilitated by birds represents the most parsimonious explanation for the phylogeographic and biogeographic patterns of organisms with limited active dispersal capacities [17–21].

The focus of our study is on *Ramulus mikado*, a predominantly parthenogenetic stick insect species found commonly in Japan, with only a few documented instances of males [22]. Notably, avian endozoochory has been demonstrated in *R. mikado* (= *R. irregulariterdentatum*) eggs in experimental conditions (Fig. 1) [12], suggesting the possibility of historical avian-mediated dispersal in natural settings. In this investigation, we analyze the population genetic structure of *R. mikado* to determine the influence of potential geographical barriers on their phylogeographic structure. We utilized a multifaceted molecular approach, incorporating mitochondrial sequences, SSR markers, and genome-wide SNPs to determine the phylogenetic relationships among individuals collected throughout the species distribution range. Mitochondrial sequence analysis with maternally inherited markers is anticipated to reveal a genetic structure similar to that of sexually reproducing species, even in predominately parthenogenetic species. SSR markers, which have high mutation rates, are probably appropriate for detecting limited intraspecific genetic variation in parthenogenetic species. Additionally, genome-wide SNP markers can offer more reliable data for estimating kinship relationships among individuals sharing a common parthenogenetic ancestor.

**Fig 1.**
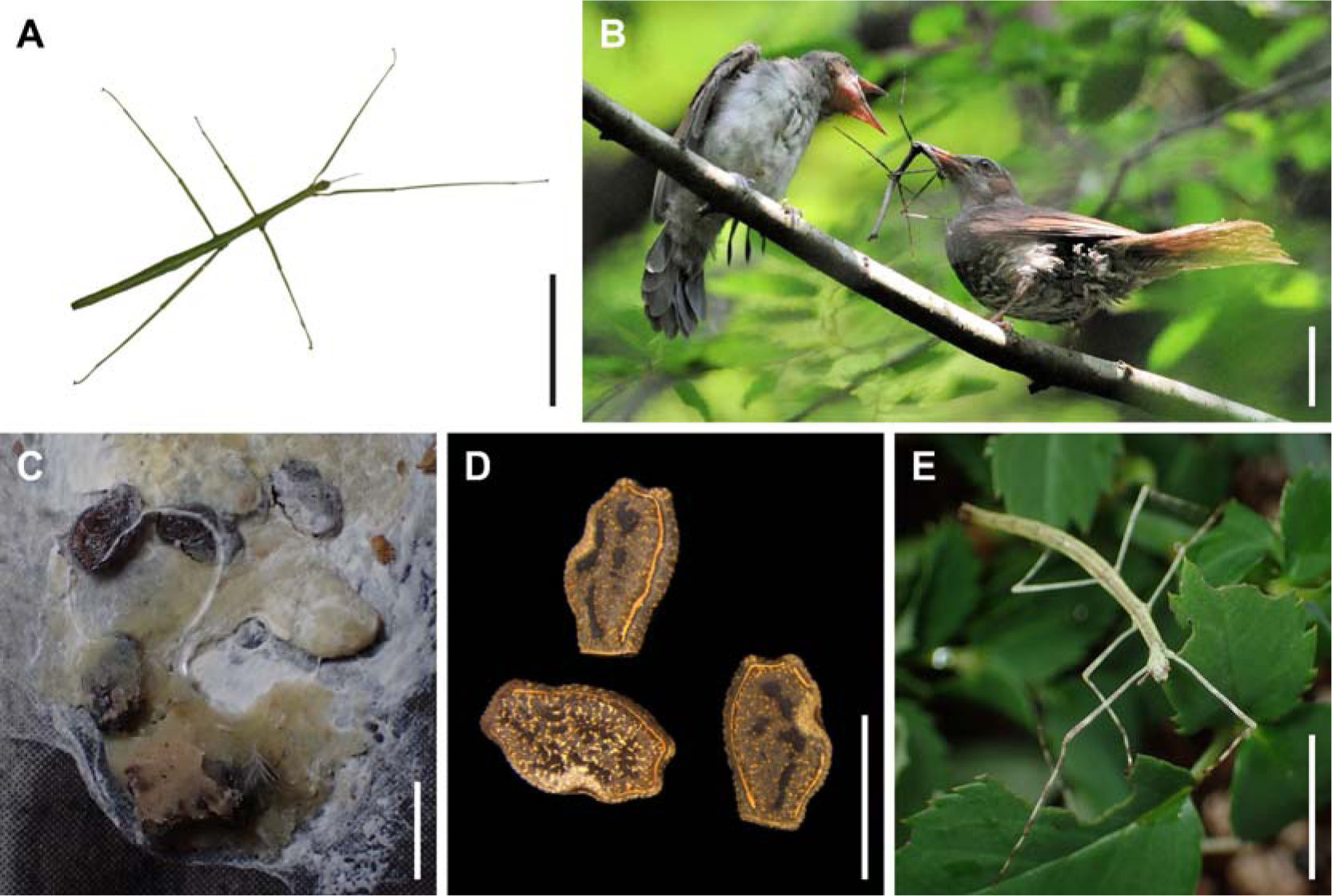
(A) The female adult of the stick insect *Ramulus mikado*. (B) The brown-eared bulbul (*Hypsipetes amaurotis*) feeding *R. mikado* to its chicks. (C) The *H. amaurotis* fecal pellets containing intact *Ramulus mikado* eggs. (D) Intact *Ramulus mikado* eggs recovered from *H. amaurotis* faeces. (E) First instar nymph of *R. mikado* hatched from the excreted egg. Scale bars: 50 mm (A), 100 mm (B), and 2 mm (C–E).

By integrating the results from these three genetic markers, each with distinct advantages, we have examined the intraspecific phylogeographic pattern of parthenogenetic and flightless stick insect species with unparalleled resolution. Consequently, we discovered unique phylogeographic patterns in the flightless stick insect, such as the identification of identical mitochondrial COI haplotypes across significant distances, which lends support to the hypothesis of passive long-distance genotypic dispersion.

## 2. Materials and Methods

### (a) Study species, sample collection, and DNA extraction

*Ramulus mikado* is a predominantly parthenogenetic and flightless stick insect widely distributed throughout Japan (Fig. 1) [22,23]. While the genus *Ramulus* encompasses over 100 species, *R. mikado* represents the sole species within this genus in Japan [22,24]. Between 2014 and 2018, we collected 67 *R. mikado* specimens from two islands in the Japanese Archipelago (Fig. 2 & Table S1). Males are rarely documented in this species, and all the specimens we collected were females. Specifically, we obtained 31 individuals from Shikoku Island and 36 from Honshu Island. Due to the extensive sampling area, we divided Honshu Island into West Honshu and East Honshu, based on the Itoigawa-Shizuoka Tectonic Line at the westernmost side of the Fossa Magna in central Honshu. The genetic structure of numerous insect groups reflects the separation of eastern and western Japan in the Fossa Magna region, where the current geological structures were formed 0.7–1.0 million years ago [25,26].

**Fig 2.**
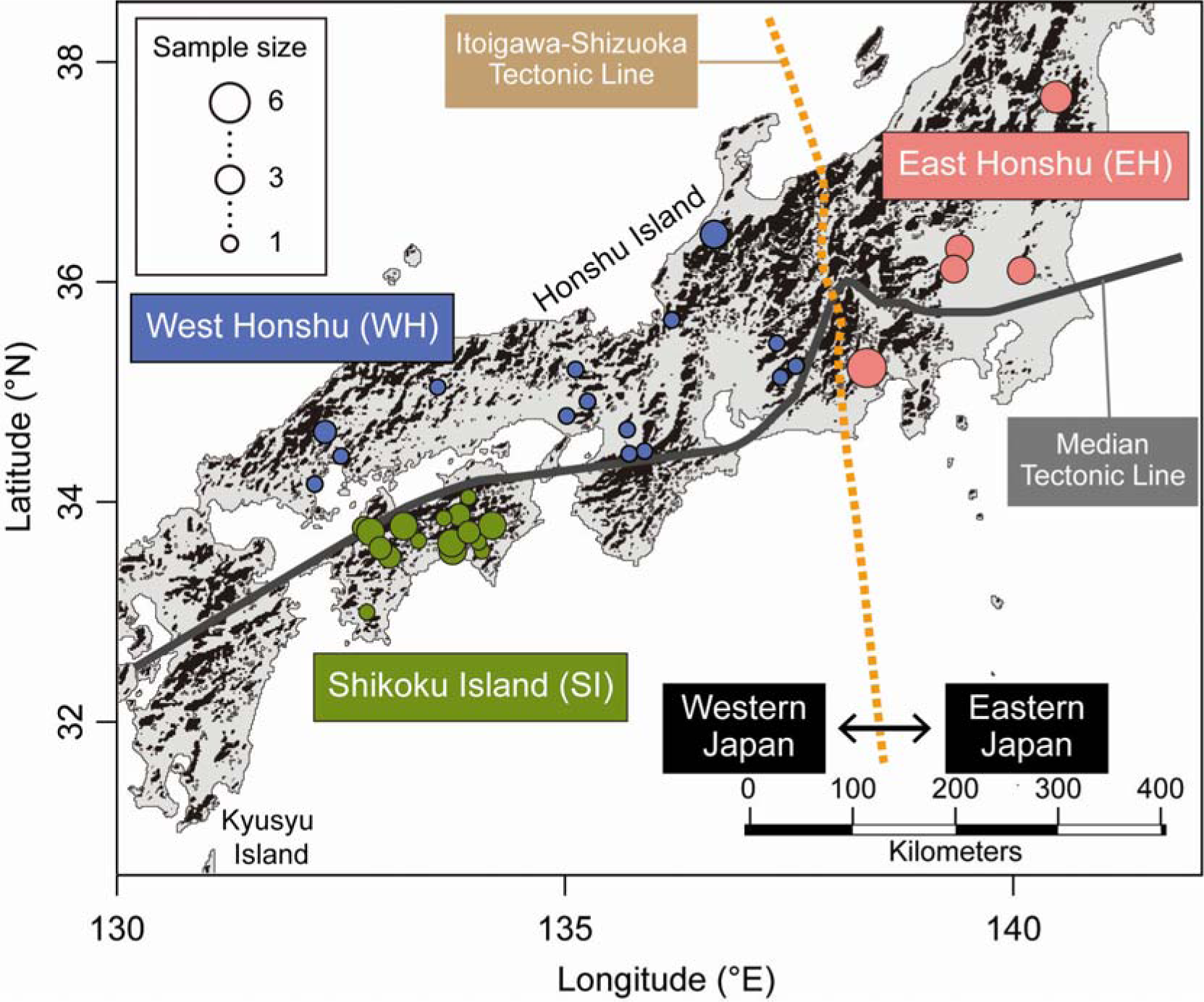
Map illustrating the sampling locations of *Ramulus mikado* used in this study. The map was generated using maptools in R software (version 3.6.2; http://cran.r-project.org/) with data sourced from the National Land Numerical Information, Ministry of Land, Infrastructure, Transport, and Tourism, Japan (https://nlftp.mlit.go.jp/index.html). Samples collected within a distance of fewer than 10 km were combined.

For data analysis, three populations were defined based on their respective regions: Shikoku Island (SI), West Honshu (WH), and East Honshu (EH). To obtain genomic DNA, *R. mikado* samples were preserved in absolute ethanol. Subsequently, the Gentra Puregene Tissue Kit (QIAGEN) was employed to extract total genomic DNA, following the manufacturer’s instructions. This extracted DNA was used for the subsequent molecular analysis.

### (b) Mitochondrial DNA analysis

For the analysis of genetic diversity, phylogenetic relationships among haplotypes, and their geographic distribution among three populations, we determined the sequences of the mitochondrial DNA cytochrome oxidase subunit I (COI) gene. The COI gene was amplified and sequenced using primers LCO_Phas3 (5’– AAC TCA GCC ATT TTA CTA ATG AAA CG –3’) and HCO_Phas1 (5’– TAT ACT TCT GGA TGA CCA AAAAAT CA –3’), which were designed based on the sequences of related taxon found in the DNA Data Bank of Japan (DDBJ). PCR amplification was conducted in a GeneAmp PCR System 2700 thermal cycler (Applied Biosystems, Foster City, California, USA). The PCR products were purified by using a High Pure PCR product purification kit (B A High Pure PCR Product Purification Kit (Roche Diagnostics, Mannheim, Germany), and purified products were sequenced directly using an ABI BigDye Terminator Cycle Sequencing Kit ver. 3.1 (Applied Biosystems) on the ABI PRISM 3130 Genetic Analyzer (Applied Biosystems). Both strands of the amplified PCR products were sequenced, and electropherograms were assembled using Finch TV (http://www.geospiza.com/finchtv/). The assembled sequences were aligned with all analyzed samples using CLUSTAL W [27] with default settings in MEGA-X [28]. We determined the mtDNA haplotype based on this aligned data.

To compare mitochondrial genetic variation among populations, we calculated the number of haplotypes, haplotype diversity, and nucleotide diversity using DnaSP 6 [29,30]. We employed analysis of molecular variance (AMOVA) to investigate genetic variance and differences among populations. Overall and pairwise *F*_ST_ values were calculated, and the probability of each pairwise *F*_ST_ value being different from zero was tested based on 999 permutations using Arlequin version 3.5 [31]. To evaluate the genetic relationships among genotypes, we constructed a neighbor-net network using SplitsTree version 4.15.1 [32] and performed principal coordinates analysis (PCoA) using GenAlEx 6.5 [33]. Neighbor-net networks were generated from COI gene sequence alignments based on uncorrected *p*-distance matrix. The genetic distance for PCoA was computed employing the Maximum Composite Likelihood model [34] with MEGA-X [28]. The association between geographic and genetic distance was evaluated for 1) all populations, 2) SI and WH populations, 3) WH and EH populations, and 4) each of the three populations using Mantel tests by GenAlEx version 6.5.

### (c) SSR analysis

We successfully isolated 13 SSR loci from the nuclear genome of *R. mikado* and designed corresponding SSR primers (Supplementary note 1). We employed these newly designed primers to determine genotypes for analyzing genetic diversity among the three populations, phylogenetic relationships among multilocus SSR genotypes, and their geographic distribution. We performed PCR amplification of the 13 SSR loci using 5 μL reactions with the QIAGEN Multiplex PCR Kit and a fluorescent dye-label protocol [35]. Each reaction contained 10 ng of genomic DNA, 2.5 μL of Multiplex PCR Master Mix, 0.01 μM of forward primer, 0.2 μM of reverse primer, and 0.1 μM of fluorescently labeled primer. The amplification protocol consisted of 95 °C for 15 min, 33 cycles at 94 °C for 30 s, 57 °C for 1.5 min, and 72 °C for 1 min, followed by an extension at 60 °C for 30 min. We determined product sizes using an ABI PRISM 3130 Genetic Analyzer and GeneMarker software (SoftGenetics, State College, Pennsylvania, USA).

We established multilocus genotypes based on 13 SSR loci genotypes. The probability of identical genotypes resulting from sexual reproduction was calculated utilizing GenAlEx 6.5 [33]. For each population, genetic diversity was assessed by evaluating the average number of alleles per locus (*A*), allelic richness (*R*_S_), observed heterozygosity (*H*_O_), expected heterozygosity (*H*_E_), and the fixation index (*F*_IS_). These parameters were calculated using GenAlEx 6.5, except for allelic richness, which was calculated using FSTAT 2.9.3 [36]. To evaluate the genetic relationships among genotypes, we constructed a neighbor-net network using SplitsTree version 4.15.1 [32] and performed principal coordinates analysis (PCoA) using GenAlEx 6.5. Nei’s genetic distance *D*_A_ [37] among multilocus genotypes calculated by POPULATION 1.2.30 [38] was used for the genetic distance of the network construction and PCoA. Additionally, the association between geographic and genetic distance *D*_A_ was evaluated for 1) all populations, 2) SI and WH populations, 3) WH and EH populations, and 4) each of the three populations through Mantel tests. Mantel tests were performed using GenAlEx.

### (d) MIG-seq analysis

MIG-seq is a recently developed genome-wide genotyping methodology employing a high-throughput sequencing platform [39] This technique is a microsatellite-associated DNA sequencing approach, a form of reduced representation sequencing that encompasses restriction site-associated DNA sequencing (RAD-seq)[39]. MIG-seq has recently emerged as a potent instrument for population genetics research [40,41]. A MIG-seq library was prepared following the protocol suggested by the development team [42] and sequenced utilizing the MiSeq system (Illumina, San Diego, CA, USA) and MiSeq Reagent Kit v3 (150 cycle). The raw genome-wide SNP data was archived in the DDBJ Sequence Read Archive (DRA, accession number DRA016238).

Upon eliminating primer sequences and low-quality reads [43], 9558552 reads (144827 ± 3256 reads per sample) were acquired from 9878722 raw reads (149678 ± 3360 reads per sample). The Stacks 2.62 pipeline was employed for de novo single nucleotide polymorphism (SNP) discovery [44], utilizing the following parameters: minimum depth of coverage required to generate a stack (*m*) = 3, maximum distance permitted between stacks (*M*) = 2, and the number of mismatches allowed between sample loci during catalog construction (*n*) = 2. SNP sites containing fewer than three minor alleles were filtered, and only SNPs retained by 33 or more samples were extracted. We restrict data analysis to only the first SNP per locus to avoid linkage between SNPs. The SNP filtering for excess heterozygosity was not performed because of the predominantly parthenogenetic reproduction of *R. mikado* [22]. Ultimately, 980 SNPs were procured for subsequent analyses. During these processes, one sample (EH09) was excluded due to its high missing rate. The nucleotide diversity (*N*_D_), observed and expected heterozygosity was assessed by the program populations of Stacks. Number of allelic differences between two individuals was calculated using R package poppr 2.9.4 [45]. Overall and pairwise *F*_ST_ values were calculated, and the probability of each pairwise *F*_ST_ value being different from zero was tested based on 999 permutations using GenAlEx 6.5 [33]. A neighbor-net network was also constructed by employing the uncorrected *p*-distance matrix and disregarding ambiguous sites, with the use of SplitsTree version 4.15.1 [32]. Moreover, to assess potential genetic structure, a principal coordinates analysis (PCoA) was performed based on the genome-wide SNPs using R package dartR 2.7.2. [46,47]. The correlation between geographic and genetic distance was assessed for 1) all populations, 2) SI and WH populations, 3) WH and EH populations, and 4) each of the three populations through Mantel tests and GenAlExversion 6.5.

## 3. Results

### (a) Genetic variation and difference among *R. mikado* populations

Genetic analysis based on mitochondrial COI sequences and nuclear SSR markers revealed a notable accumulation of mutations in *R. mikado* due to parthenogenetic reproduction. The observed number of alleles at the 13 newly developed loci ranged from 1 to 20 (Table S3). The observed heterozygosity for eight loci was either 0 or nearly 0, while the remaining five loci displayed high values ranging from 0.64 to 1.00. From the 67 samples, 55 multi-locus genotypes were identified. The combined random match probability for the 13 loci was 5.53 × 10^-6^, with a high power for discriminating among individuals. The MIG-seg based SNP heterozygosity (the highly heterozygous and homozygous loci) corresponds to these SSR heterozygosity pattern (Fig. S2).

In the mitochondrial COI region sequences, 39 haplotypes were identified, among which 11 haplotypes were present in multiple individuals (Table 1). The number of individuals sharing the same haplotype ranged from 2 to 6, and no specific haplotype was predominantly distributed. Intriguingly, some individuals collected from distant sites exhibited the same haplotype, and eight haplotypes were distributed in sites more than 10 km apart (Fig. S1; Table 1). The farthest Hap04 was confirmed from 683 km, and Hap06 was confirmed from 452 km. In the nuclear SSR genotype, the number of individuals exhibiting identical multi-locus genotypes was small, with only 2 or 3 observed, and no widespread specific genotype was observed. For Gen34 and Gen30 genotypes, they were collected from 19 km and 10 km away, respectively (Table 1). Two individuals exhibiting the Gen34 genotype also shared the same mitochondrial haplotype. Although the SNP analysis did not obtain the same genotype, individuals with close genetic distances across distant populations have been confirmed. The number of allelic differences between two individuals ranged from 40 (2.04%) to 136 (6.94%). The minimum number of allelic differences between two individuals was observed between EH13 and WH03, which were 259 km apart. The values of haplotype diversity *H*_D_ and the nucleotide diversity of COI sequences, as well as the allelic richness *R*_S_ of SSR markers, which are crucial indicators for assessing genetic diversity, were relatively high in the SI and WH populations and low in the EH population (Table 2). Nonetheless, no significant differences were observed in the nucleotide diversity of genome-wide SNPs among the populations. AMOVA analysis based on COI haplotypes and SSR genotypes revealed significant genetic differences among the three populations (Table 3). The *F*_ST_ values based on mitochondrial sequence data, SSR genotype data as well as genome-wide SNP data were all significant among the populations (p < 0.01). Each *F*_ST_ value calculated from mitochondrial sequence data and genome-wide SNP data was significant for every pair of populations, while the pairwise *F*_ST_ value between the EH and WH populations did not exhibit a significant difference when assessed using SSR genotype data (p = 0.218; Table 3).

**Table 1.** Mitochondrial DNA haplotype and nuclear SSR genotype detected from multiple samples.

|  |  |  |  |
| --- | --- | --- | --- |
| Haplotype or Genotype | *n* | Max. Dist. (km) | Sample ID |
| Mitochondrial DNA |  |  |  |
| Hap04 | 4 | 683 | WH12, EH10, EH11, EH12 |
| Hap06 | 3 | 452 | SI26, SI27, WH20 |
| Hap01 | 5 | 179 | EH01, EH02, EH03, EH05, EH06 |
| Hap05 | 6 | 76 | WH19, EH13, EH14, EH15, EH16, EH17, EH18 |
| Hap25 | 5 | 52 | SI07, SI09, SI16, SI17, SI18 |
| Hap34 | 3 | 35 | SI22, SI23, SI24 |
| Hap32 | 2 | 18 | SI19, SI21 |
| Hap20 | 2 | 16 | SI01, SI03 |
| Hap09 | 3 | < 1 | WH22, WH23, WH24 |
| Hap03 | 3 | < 1 | EH07, EH08, EH09 |
| Hap31 | 2 | < 1 | SI14, SI15 |
| Nuclear SSR |  |  |  |
| Gen34 | 2 | 19 | SI09, SI17 |
| Gen30 | 2 | 10 | SI04, SI05 |
| Gen01 | 3 | < 1 | EH01, EH02, EH03 |
| Gen04 | 3 | < 1 | EH07, EH08, EH09 |
| Gen07 | 3 | < 1 | EH13, EH16, EH17 |
| Gen11 | 2 | < 1 | WH22, WH23, WH24 |
| Gen03 | 2 | < 1 | EH05, EH06 |
| Gen06 | 2 | < 1 | EH11, EH12 |
| Max. Dist: Maximum distance between individuals with the identical haplotype or genotype |  |  |  |

**Table 2.** Genetic diversity of COI mitochondrial sequence, nuclear SSR loci and genome-wide SNP of three populations.

| Population | Mitochondrial sequence |  |  | Nuclear SSR |  |  |  | Genome-wide SNP |  |  |
| --- | --- | --- | --- | --- | --- | --- | --- | --- | --- | --- |
| | $N_H$ | $H_D$ | $N_D$ | $A$ | $R_S$ | $H_0$ | $H_E$ | $N_D$ | $H_0$ | $H_E$ |
| ALL | 39 | 0.972 | 0.0077 ± 0.0005 | 3.0 | 3.5 | 0.34 | 0.32 | 0.00135 ± 0.00005 | 0.670 | 0.393 |
| SI ($n = 31$) | 20 | 0.961 | 0.0091 ± 0.0006 | 3.9 | 3.4 | 0.37 | 0.32 | 0.00136 ± 0.00005 | 0.667 | 0.390 |
| WH ($n = 18$) | 16 | 0.980 | 0.0075 ± 0.0008 | 3.4 | 3.4 | 0.34 | 0.35 | 0.00136 ± 0.00005 | 0.669 | 0.385 |
| EH ($n = 18$) | 5 | 0.797 | 0.0035 ± 0.0004 | 2.4 | 2.3 | 0.30 | 0.32 | 0.00139 ± 0.00005 | 0.677 | 0.390 |

$N_H$ : Number of haplotype, $H_D$ : Haplotype diversity, $N_D$ : Nucleotide diversity, $A$ : Number of alleles, $R_S$ : Alleric richness, $H_0$ : Observed heterozygosity, $H_E$ : Expected heterozygosity

**Table 3.**
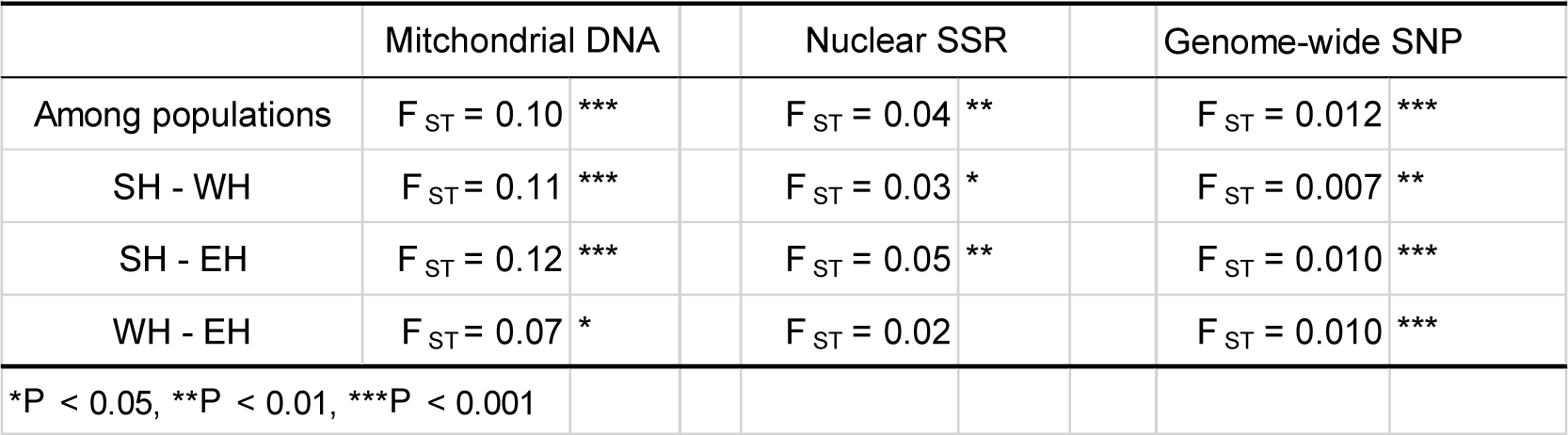
Genetic differentiation among populations and pairwise population comparison.

|  |  |  |  |  |  |  |  |  |
| --- | --- | --- | --- | --- | --- | --- | --- | --- |
|  | Mitochondrial DNA |  |  | Nuclear SSR |  |  | Genome-wide SNP |  |
| Among populations | $F_{ST} = 0.10$ | *** | | $F_{ST} = 0.04$ | ** | | $F_{ST} = 0.012$ | *** |
| SH - WH | $F_{ST} = 0.11$ | *** | | $F_{ST} = 0.03$ | * | | $F_{ST} = 0.007$ | ** |
| SH - EH | $F_{ST} = 0.12$ | *** | | $F_{ST} = 0.05$ | ** | | $F_{ST} = 0.010$ | *** |
| WH - EH | $F_{ST} = 0.07$ | * | | $F_{ST} = 0.02$ | | | $F_{ST} = 0.010$ | *** |
| *$P < 0.05$, **$P < 0.01$, ***$P < 0.001$ | | | | | | | | |

### (b) Genetic relationship among individuals and their spatial distribution

Our molecular analyses, which employed COI haplotypes, SSR markers, and MIG-seq analysis, revealed limited associations between lineage and geographic distribution (Fig. 3–5), with some exceptions, such as COI haplotypes and genome-wide SNPs consisting exclusively of samples from the SI population (Fig. 3A, C and 4A, C). However, aside from these few instances, the correspondence between phylogenetic relationships and distribution was not evident. In all analyses of genetic markers, individuals with close genetic relationships were found to be dispersed across different populations. This pattern remained consistent across distinct methods employed in phylogenetic analysis and measures of genetic distance (Data not presented here).

**Fig 3.**
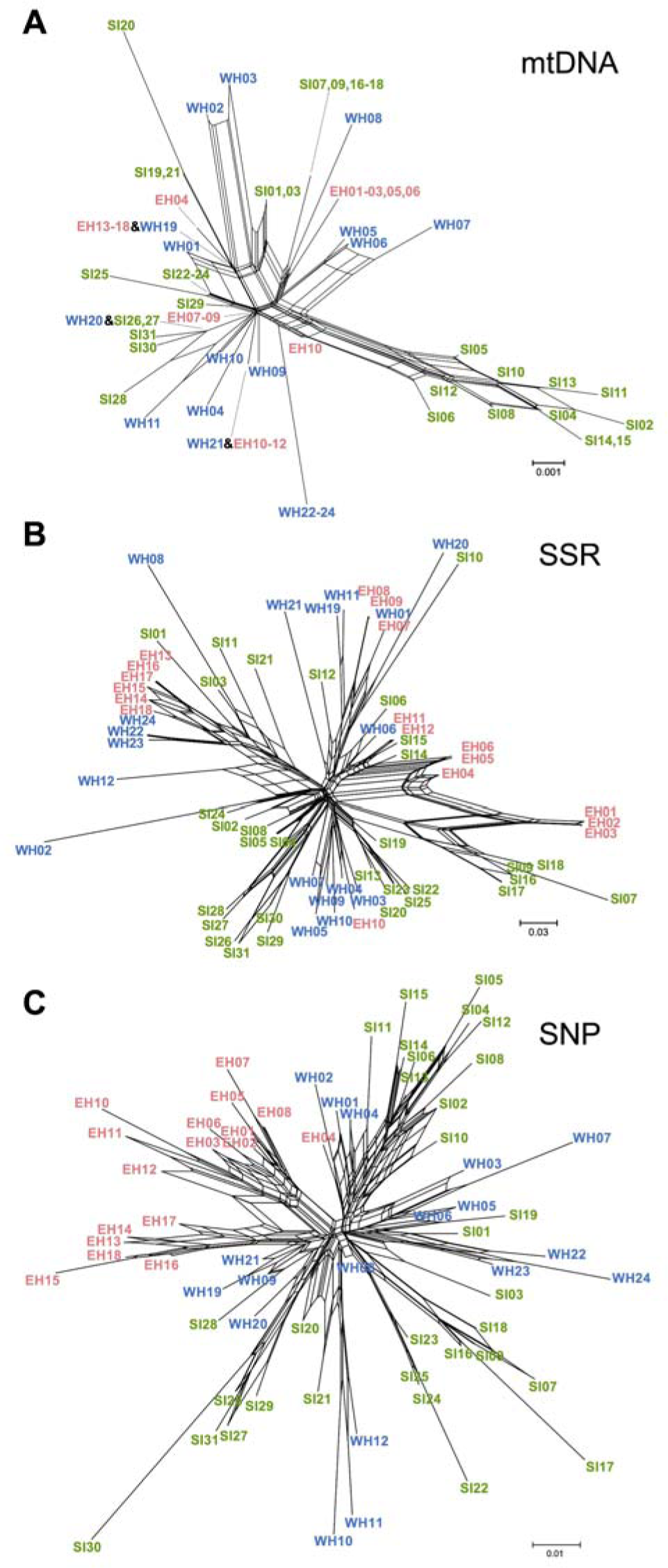
Neighbor-Net network of *Ramulus mikado* samples reconstructed based on the COI haplotypes (A), nuclear SSR (B), and genome-wide SNP (C). Branch length denotes the average number of substitutions per site.

**Fig 4.**
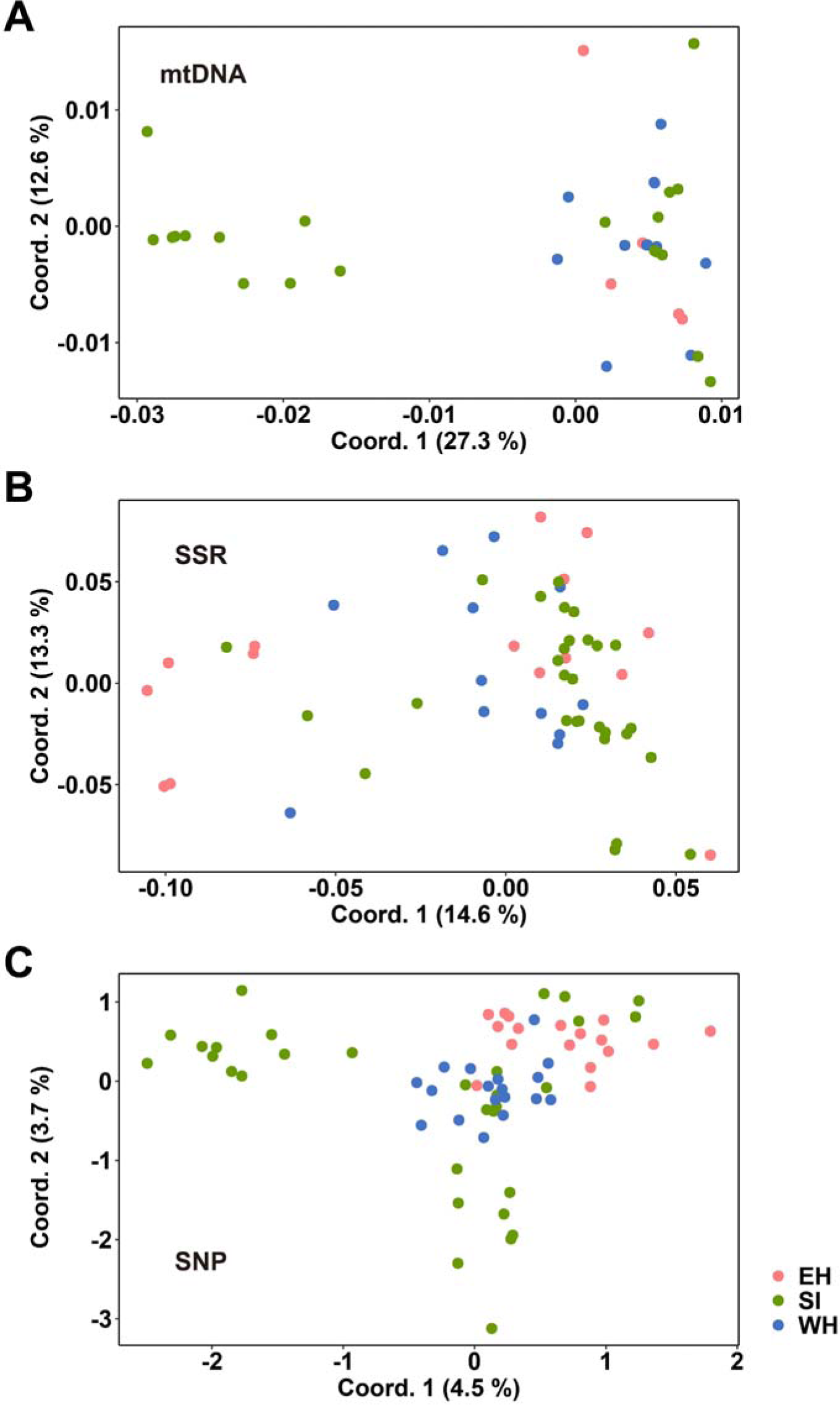
Principal coordinate analysis plot based on COI haplotypes (A), nuclear SSR (B), and genome-wide SNP (C) data.

**Fig 5.**
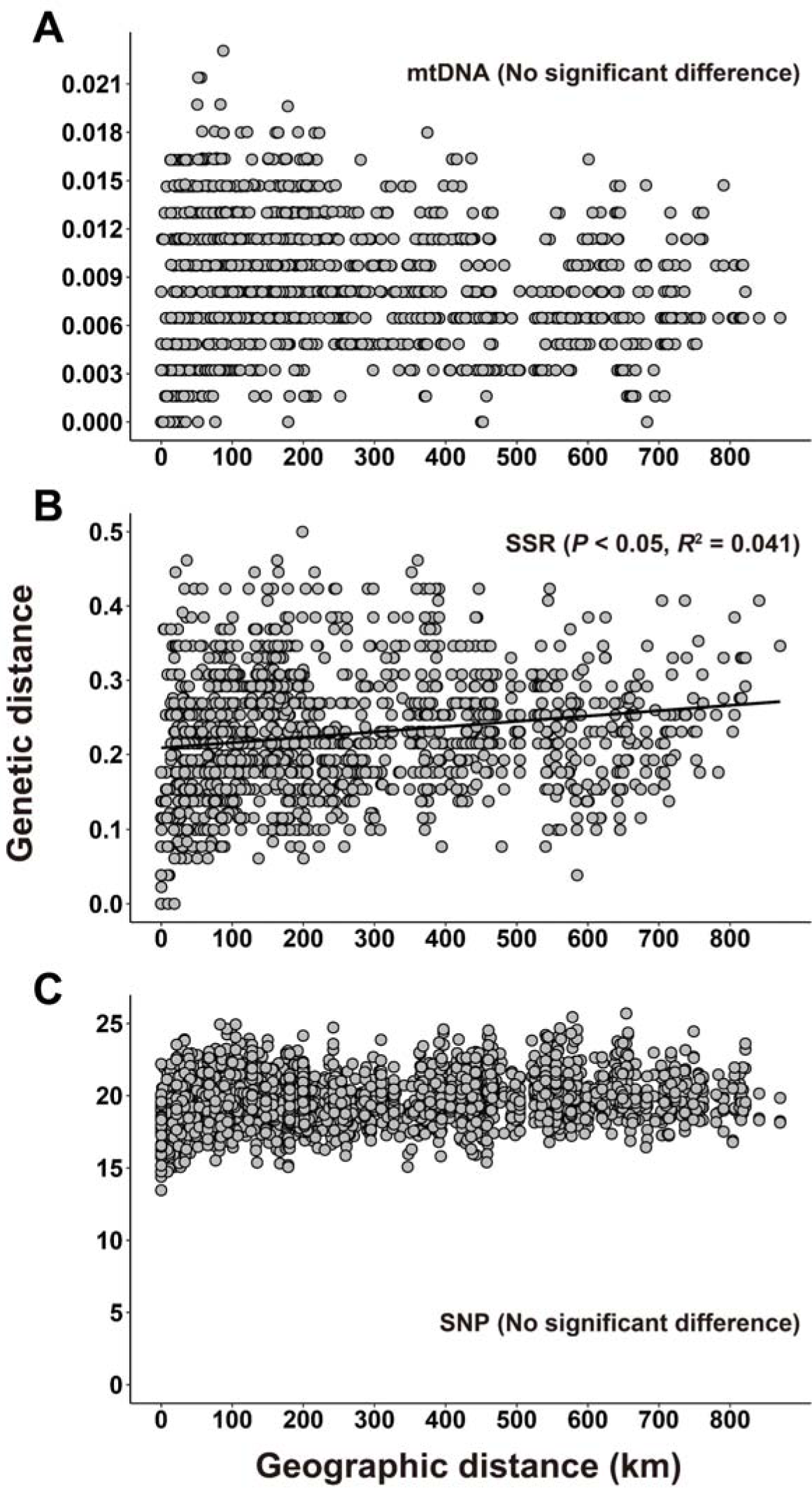
Relationships between geographic and genetic distances at the individual level based on COI haplotypes (A), nuclear SSR (B), and genome-wide SNP (C) data.

The Mantel test results, examining the correlation between genetic distance and geographical distance within specific regions, yielded different outcomes among regions (Fig. 5). The Mantel test, using all samples, revealed no significant correlation between geographical distance and genetic distance in the mitochondrial sequence and genome-wide SNP data (Fig. 5A, C). Although a weak yet significant correlation was detected in the SSR data (p < 0.05, Fig. 5B), the R-squared value was low (*R*^2^ = 0.041). Notably, genetically very similar individuals are found up to approximately 700 km in the mitochondrial sequence data and up to about 600 km in the SSR data. The combinations of individuals with the highest level of genetic distance between individuals were observed even at a distance of about 100 km in all three genetic markers. Within the samples merging Western Honshu (WH) and Shikoku Island (SI), “isolation by distance” was not observed based on COI haplotypes, SSR and SNP data (Fig. S3). In contrast, within Eastern Honshu (EH), genetic distance exhibited a significant positive correlation with geographical distance based on all molecular methods. This correlation was particularly pronounced in SSR markers (Fig. S3, p < 0.05, *R*^2^= 0.678).

## 4. Discussion

Although the majority of research on avian-mediated dispersal has primarily focused on seed dispersal, birds can disperse a diverse range of invertebrates via endozoochory. Extreme cases include the transportation of insect eggs or larvae, as well as fish eggs, potentially facilitating the colonization of novel habitats [12,48–50]. It is essential to note, however, that these studies demonstrate passive egg dispersal under laboratory conditions. Our multifaceted molecular analysis, covering different time scales suggests that long-distance dispersal, likely through avian predation, can substantially impact the distribution and population structure of *R. mikado* (Fig. 3–5, S1 & Table 1). These insights highlight the potential role of birds in shaping ecosystems and emphasize the need for further investigation into the ecological consequences of avian-mediated dispersal.

Our SSR analysis shows high heterozygosity in five loci and zero or near-zero in the remaining eight loci in *R. mikado* (Table S3). This pattern, also observed in genome-wide SNP data (Fig. S2), suggests the predominance of asexual reproduction, though historical cryptic gene flow cannot be excluded [51]. This assumption is consistent with not only the female predominance in *R. mikado* [22] but also the non-functionality of the rare males (Nozaki et al., unpublished data). The pattern of heterozygosity contrasts with terminal fusion automixis or gamete duplication, causing tremendous heterozygosity loss [52,53]. Automixis with central fusion seems most plausible, as SSR analysis revealed mothers and offspring (embryos) mostly shared genotypes, with rare heterozygous to homozygous transitions due to recombination (Nozaki et al., unpublished data). The mixture of highly heterozygous and homozygous loci likely results from varying recombination probabilities by loci [53].

We identified some differences in COI haplotype frequency and SSR allele distribution among *R. mikado* individuals (Tables 2 and 3). In parthenogenetic species, typically a single strain or few strains with limited genetic variation have widespread distribution due to efficient parthenogenic reproduction and rapid expansion in a short evolutionary time scale [54,55]. The accumulation of genetic variation implies a relatively long parthenogenetic persistence, allowing mutation accumulation. Given insect mitochondrial DNA substitution rates, including the COI region, range from 1.5% [56] to 2.3 % [57] per million years, the 0.77% nucleotide diversity of COI sequences likely reflects a history of 0.34–0.51 million years. Although multiple parthenogenetic origins could account for these differences, SSR marker or genome-wide phylogeny supports a single lineage radiation pattern, inferring a single parthenogenesis origin in *R. mikado*.

Although *R. mikado* exhibits a certain degree of genetic variation, only a weak geographic signal was detected among *R. mikado* individuals with limited active dispersal ability. Notably, no positive correlation between geographical and genetic distances was observed in SI and WH populations, suggesting genotypic dispersal within these sea-separated populations. The detection of COI haplotypes Hap04 and Hap06 at distances of 680 km and 450 km, respectively, implies rapid expansion of these strains over hundreds of kilometers across the Fossa Magna, whose current geological structures had already formed before the origin of *R. mikado* (0.7–1.0 million years ago vs. 0.34–0.51 million years ago) [25,26], outpacing mutation. The PCoA plots as well as Neighbor-Net networks, based on COI and genome-wide SNP data have also revealed some distinct area-specific lineages in the SI population, while other SI lineages mixed with WH and EH populations, suggesting that some genotypes suffer long-distance dispersal. Nonetheless, these patterns also suggest that long-distance dispersal events are infrequent, as the genetic structure between populations remains partially intact, and some regional genetic differentiation is still discernible.

Distinct patterns in the EH population, which is likely a recently formed population following the last glacial period, may also support the infrequency of long-range dispersal. Pollen analysis indicates that coniferous forests predominantly covered this region during the last glacial period, while temperate broadleaf forests, which serve as suitable habitats for *R. mikado*, did not expand until after the last glacial period [58]. The low genetic variation of mitochondrial DNA and nuclear SSR markers in the EH population (Table 2) likely reflects the relatively brief history of this population and the limited number of founders that have arrived from WH and SI populations. Notably, a distinct positive correlation between geographic and genetic distance was observed in the EH population based on SSR markers (Fig. S3). The isolation-by-distance pattern in the EH population may be attributed to the limitation of long-distance dispersal and the higher mutation rate of SSR markers. Consequently, the genetic structure of the EH population is likely attributable to a relatively brief history of distribution expansion and more rapid accumulation of mutations in SSR loci compared to the rate of distribution expansion.

As discussed earlier, the unexpected phylogeographic pattern in the flightless stick insect is likely facilitated by long-distance dispersal. Avian predation emerges as the most plausible explanation for the dispersal, especially given the demonstrated resilience of their eggs in withstanding passage through avian digestive tracts [12]. Although other animals, such as mammals, could potentially disperse the eggs, birds are the most probable dispersers at distances spanning more than several kilometers, since most other animals, including mammals, typically disperse over tens or hundreds of meters [21,59]. Additionally, despite the seed-like appearance of the eggs, the possibility of egg dispersal via granivorous birds seems improbable, as granivores have evolved to crush seeds in their gizzards, leading to complete digestion of the eggs [60]. Consequently, the predation of gravid females with eggs represents the most plausible avian internal dispersal mechanism [12].

Oceanic dispersal has also contributed to the phylogenetic pattern of stick insects, as certain phasmid eggs, such as those of the Mascarene stick insects, adhere to branches or leaf surfaces and can be rafted, allowing for transport across oceanic currents [61]. Additionally, some phasmids, like *Megacrania*, possess sponge-like eggs that can float for long periods without needing to be attached to vegetation [14]. However, these dispersal methods are improbable in *R. mikado* since their eggs do not possess either a sponge-like structure or the ability to stick to the branches. While anthropogenic translocation represents a potential explanation [54,62,63], *R. mikado* specimens with identical COI haplotypes, distinguished by relatively slow mutation rates, nearly always display some variation in SSR and MIG-seq genotypes (excepting SH-09 and SH-17 in SSR genotypes), which exhibit faster mutation speeds. In addition, if anthropogenic translocation plays a role in shaping the genetic structure of this species, a similar genetic structure observed in the SI and WH populations should also be present in the EH population. Consequently, the probability of recent unintentional anthropogenic translocation, such as the transportation along with tree saplings, is exceedingly low. Instead, it is more likely that long-distance dispersal resulted from historical events that occurred over a sufficient duration to allow for the accumulation of differences in SSR and MIG-seq genotypes.

Overall, we have identified distinct phylogeographic patterns, including the identical COI genotypes covering significant distances. This supports the idea of passive long-distance dispersal of the genotypes. The question of how organisms with limited active dispersal capabilities achieve extensive distribution has captured curiosity since the time of Darwin [64]. Given that *R. mikado* possesses limited active dispersal abilities, avian internal dispersal probably aids in establishing populations across vast geographical areas. Based on the phylogeographic pattern in conjunction with prior experimental evidence [12], we propose that *R. mikado* eggs can occasionally survive gut passage in the wild, thus facilitating long-distance dispersal. These studies offer novel insights as they challenge the long-standing belief that insects inevitably perish upon avian predation. Given the capacity of parthenogenetic reproduction to facilitate avian internal dispersal of eggs within gravid female stick insects and to enhance the likelihood of successful colonization after dispersal [12], similar passive dispersal and subsequent distributional expansion might be prevalent among parthenogenetic species. Further investigation into the extent of this phenomenon is warranted.

## Data accessibility

The mitochondrial COI, SSR loci, and genome-wide SNP data are available in the DNA Data Bank of Japan (DDBJ) Sequence Read Archive (DRA; https://www.ddbj.nig.ac.jp/dra/index-e.html): LC767280–LC767346, LC623838–LC623855, and DRA016238, respectively.

## Authors’ contributions

K.S. conceived and designed the study with input from S.K. K.S., T.N., and S.F. collected samples. S.K., K.S., S.K.H., and S.F. conducted molecular experiments. K.I. and Y.S. supervised the experiments conducted by S.F. and S.K.H., respectively. K.S., S.K.H., and S.K. curated and analyzed the data, and validated the findings. K.S., T.N., S.K.H., and S.K. visualized the results. K.S. and S.K. wrote the original draft with input from T.N., and S.K.H. All authors revised the manuscript and approved the final version.

## Acknowledgments

The authors thank Junpei Haga, Katsumi Iwahori, Kota Sakagami, Mineki Yoshida, Naoyuki Nakahama, Osamu Tominaga, Seiki Honda, Shumpei Kitamura, and Yuta Mashimo for providing specimens. We are also grateful to Koya Shishido, Kazuki Kurita, Riku Shina, Yuya Nakazawa, Ryuta Sato, Takako Shizuka, Hidehito Okada, Kazuma Takizawa, Hakuren Kato and Toshihito Takagi for their technical assistance. We also extend our appreciation to Drs. Yoshiaki Tsuda, Takeshi Yokoyama, Tatsuya Fukuda and Osamu Miura for insightful discussions.

---

**Fig S1.**
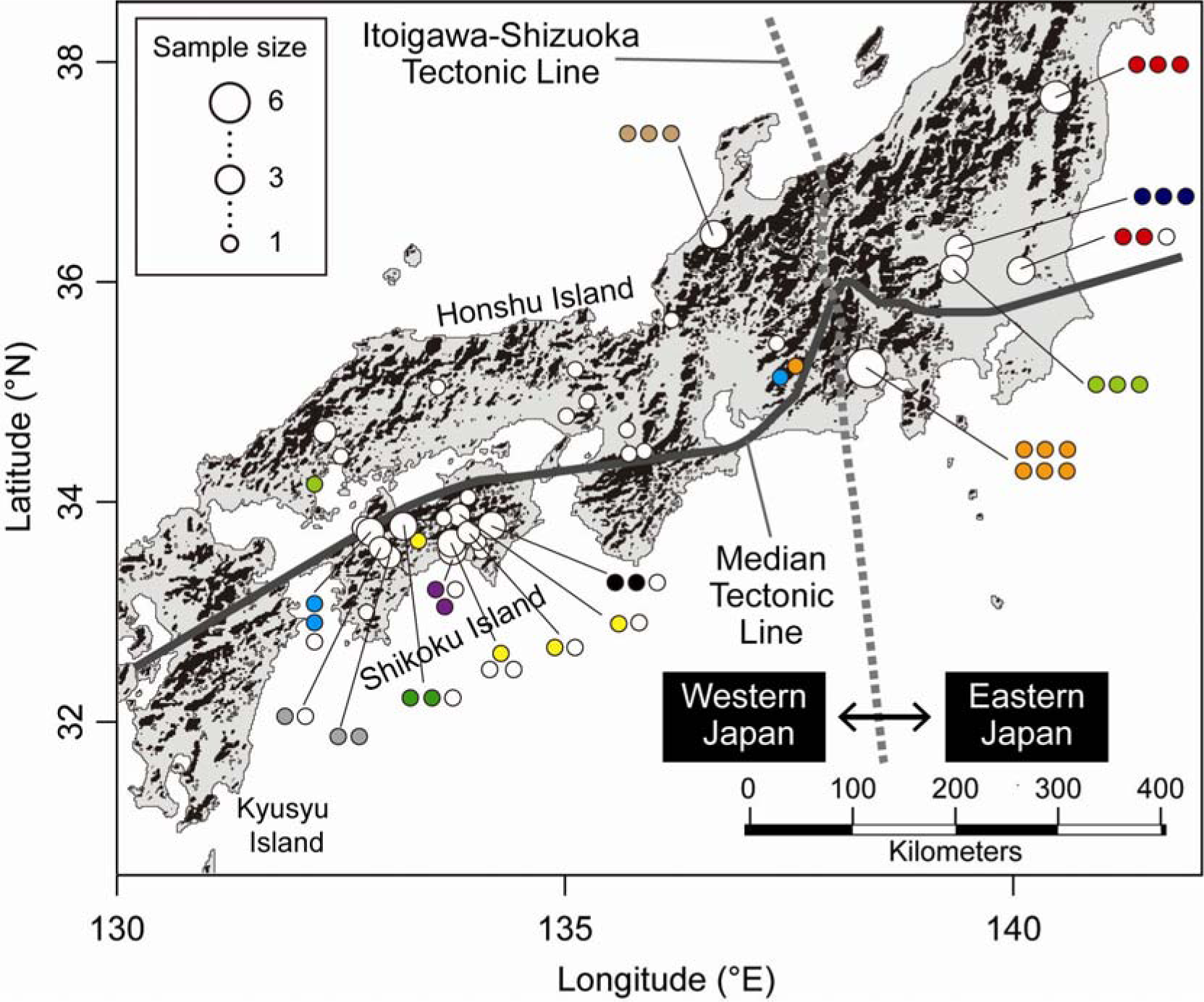
Map illustrating the COI haplotype distribution of *Ramulus mikado* within each sampling location. The map was generated using maptools in R software (version 3.6.2; http://cran.r-project.org/) with data sourced from the National Land Numerical Information, Ministry of Land, Infrastructure, Transport, and Tourism, Japan (https://nlftp.mlit.go.jp/index.html). Samples collected within a distance of fewer than 10 km were combined. Only haplotypes shared by two or more individuals are color-coded.

**Fig S2.**
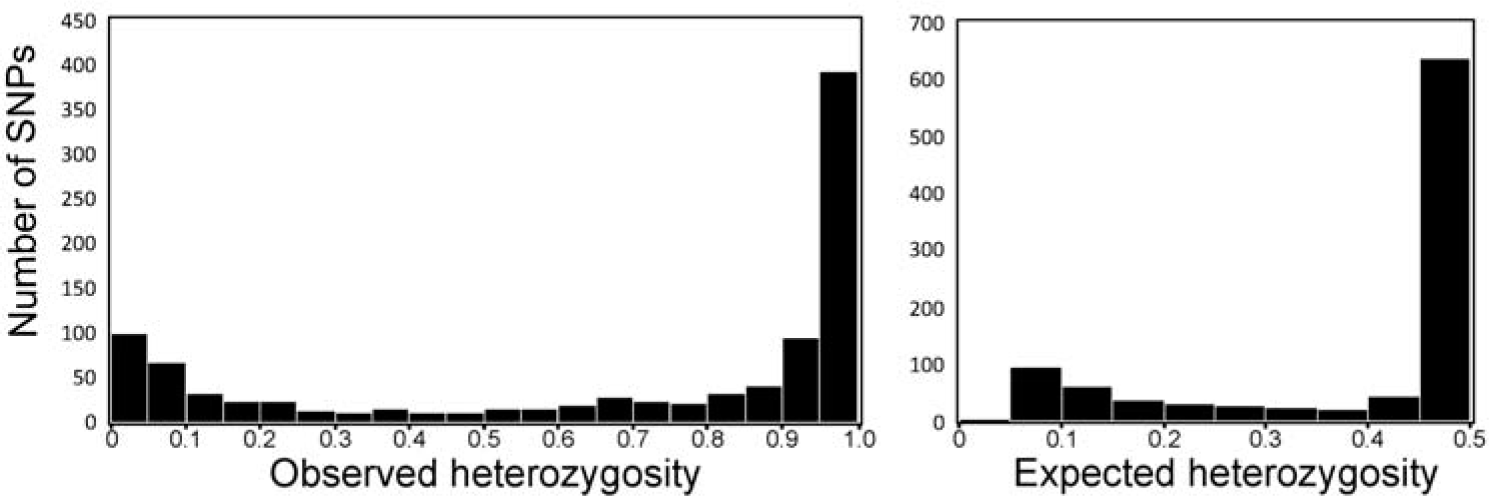
The observed and expected heterozygosity of *Ramulus mikado* based on genome-wide SNPs.

**Fig S3.**
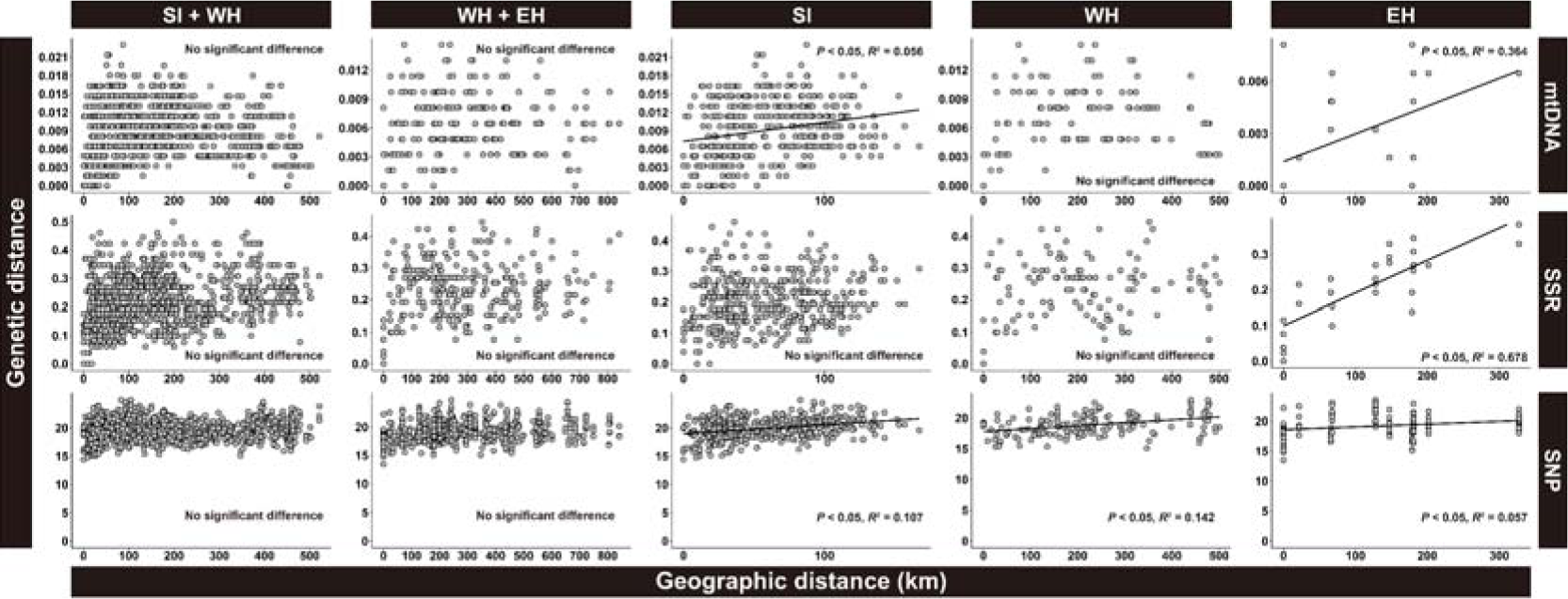
Relationships between geographic and genetic distances at the individual level based on COI haplotypes (A), nuclear SSR (B), and genome-wide SNP (C) data.

