## Supplemental note for "Phylogeographical evidence for historical long-distance dispersal in the flightless stick insect *Ramulus mikado*"

Supplementary note 1.

**Development of microsatellite markers**

DNA fragment libraries for two species were constructed using the Ion Xpress Plus Fragment Library Kit (Thermo Fisher Scientific), amplified using the Ion PGM Template OT2 400 Kit (Thermo Fisher Scientific), and then sequenced using the Ion PGM Sequencing 400 Kit (Thermo Fisher Scientific) and an Ion 318 Chip v2 (Thermo Fisher Scientific). After filtering for identical reads, 213,527 were identified for *Ramulus mikado*. These sequences were screened for potential microsatellite loci using MSATCOMMANDER [1]. Primers were designed for all sequences containing more than ten dinucleotide or eight trinucleotide tandem repeats using Primer3 software [2] with the default settings.

A total of 99 primer pairs were obtained for screening. PCR amplification was performed in 5-μl reactions using the QIAGEN Multiplex PCR Kit (QIAGEN) and a protocol for fluorescent dye-labeled primers [3]. Each reaction contained the following components: 10 ng of genomic DNA, 2.5 μl of Multiplex PCR Master Mix, 0.01 μM forward primer, 0.2 μM reverse primer, and 0.1 μM fluorescently labeled primer. Amplifications used the following setting: 95 °C for 15 min; 33 cycles at 94 °C for 30 s, 57 °C for 1.5 min and 72 °C for 1 min; and an extension at 60 °C for 30 min. Product sizes were determined using an ABI PRISM 3130 Genetic Analyzer and GeneMapper software (Applied Biosystems, Foster City, CA, USA).

After an amplification trial using 16 samples, 13 primer pairs (Table S2) showing clear peak patterns were selected for the microsatellite analysis of present study.
